# The oral microbiome and its effect on exhaled breath volatile analysis – the elephant in the room

**DOI:** 10.1101/2025.04.30.651103

**Authors:** Lorenzo S. Petralia, Anesu Chawaguta, Veronika Ruzsanyi, Chris A. Mayhew, Daniel Sanders

## Abstract

The rapid transfer of volatiles from alveolar blood into the lungs and then out of the body in exhaled breath leads to the common and natural conclusion that these volatiles provide information on health and metabolic processes, with considerable potential as biomarkers for use in the screening, diagnosis and monitoring of diseases. Whilst these exhaled volatiles could well serve as biomarkers for human metabolic processes, thereby providing insights into the clinical and nutritional status of individuals, there exist various confounding factors that limit their easy application. A major confounding factor is the introduction of microbially produced oral volatiles into the exhaled breath, yet these volatiles are often ignored in discovery volatile research studies. Here, we provide a comparative cross-sectional study of selected volatiles commonly found in exhaled breath, namely 1-propanol, 2-propanol, ethanol, acetoin, acetone, isoprene, methanol and 2-pentanone, measured in nasal and oral end-tidal exhaled breath samples for twenty-one volunteers, using the analytical technique of gas chromatography ion mobility spectrometry. Significant differences in 1-propanol, 2-propanol, ethanol and acetoin concentrations are found between breath samples exhaled via the mouth and those exhaled via the nose, serving to illustrate the extent to which the oral microbiome can influence volatile concentrations in breath. A central finding is that the nasally sampled volatiles are little influenced by the inhalation route (oral or nasal). The evidence is clear that in order to reduce the influence of the oral microbiome on untargeted discovery breath research studies, end-tidal exhaled nasal breath samples should be taken for endogenous volatile analysis, otherwise oral microbial volatiles could be falsely identified as biomarkers. This is particularly important given the rise in the use of machine learning algorithms and artificial intelligence to identify variations in volatilomes. The development and commercialisation of simple, user-friendly and comfortable end-tidal exhaled nasal sample collection devices are required for nasal sampling to become widely adopted.

**Graphical abstract / ToC figure:** 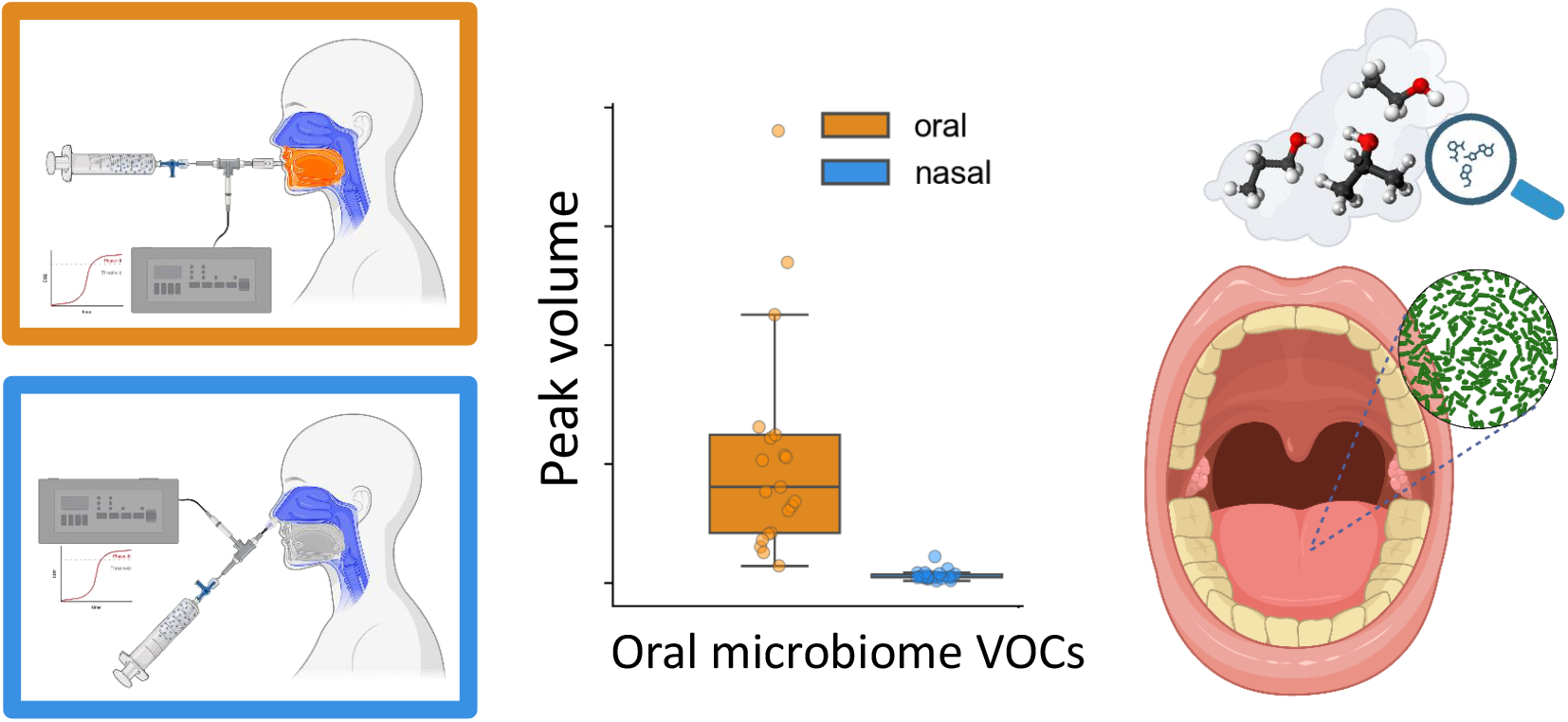

## 1. Introduction

Volatile organic compounds (VOCs) are found in various human biological matrices, including breath, faeces, saliva, skin, and urine [1–3]. However, exhaled breath is distinguished by its virtually unlimited availability and the frequency at which it can be collected, making it a practical, cost-effective and accessible biological sample with high patient acceptance [4,5]. Since VOCs in the human body travel from organs and tissue through the blood stream to the lungs, where they are then exchanged into exhaled breath, analytical techniques have detected a wide range of these compounds in exhaled breath generated by various metabolic, physiological, and pathological processes within the body [6,7]. These VOCs encompass both endogenous compounds, originating from the natural microbiome and physiological and pathological mechanisms within the body, and exogenous compounds, which are determined by external factors such as environmental, dietary and medication. The *breathprint or volatilome*, i.e., the volatile composition of an individual’s exhaled breath, can thus serve as a valuable tool for investigating pathophysiological and metabolic processes within the body, thereby making it attractive for biomedical research and potentially for clinical applications as a screening or diagnostic tool and for precision or personalised medicine [7–11].

In recent years, researchers have tried to ascertain an association between certain VOCs and specific medical conditions, with goals for early diagnostics, disease monitoring, or even predicting medical conditions [12]. Endogenous volatile compounds that potentially carry information relevant to diseased states are present in exhaled breath at low levels, typically in parts per billion by volume (ppb_v_) or less. Identifying and quantifying these low-level compounds present significant analytical, analysis and data interpretation challenges. Other problems arise from the fact that endogenous volatiles are not necessarily specific to any given disease and may not be able to detect diseases at their early stages. These challenges are compounded by potential interference and distortion from exogenous compounds that can also be present in exhaled breath at such low concentrations [13].

Further complications for breath analysis arise when analysing orally exhaled breath due to bacterial bio-transformation of substrates to volatiles within the oral cavity [14]. This poses a significant challenge for researchers aiming to diagnose systemic or metabolic conditions through exhaled breath volatile analysis, as the origins of many detected compounds remain largely unknown. Furthermore, where possible origins are identified, causations often remain vague or impossible, since most breath volatiles are typically simple chemical structures with low information content, and hence might result from various metabolic processes. An exhaustive overview of the major VOCs detected in human exhaled breath that has been compiled by Sharma et al elucidates the potential biochemical pathways involved in their generation and, in addition, discusses the diagnostic significance of their analysis and reviews the analytical techniques employed in breath testing [15].

Limited progress has been done in terms of standardization of exhaled breath sampling techniques and identifying and addressing sources of contamination and confounding factors due, for examples, to the environment (e.g., the impact of inhaled exogenous VOCs [13]) or to the microbiome [14]. With regards to the microbiome, a crucial aspect remains the choice of the exhaled breath sampling route, i.e., opting for oral or nasal exhaled breath collection. The few studies that have explored nasal versus oral breath samples point out that some volatiles observed in oral breath samples predominantly come from bacterial activity within the oral cavity [16–18]; notably, Španěl et al proved that ammonia observed in exhaled breath originates mainly in the mouth [18]. Considering that ammonia as a systemic VOC can constitute an important biomarker for kidney failure [19] (and potentially for liver dysfunction and respiratory tract infections [12,20]), the higher levels of ammonia generated in the oral cavity limit the development of any breath test for detecting/monitoring such diseases. These previous studies and their findings naturally raise the question of why orally exhaled sampling is currently almost the norm within the breath research community, despite the obvious oral cavity microbial volatile contamination issue. This matter is not sufficiently emphasised, with more focus being evidently placed on VOC discovery, analytical techniques, standardisation, complex statistical analysis and sampling devices, but not on the adopted exhaled breath sampling route. Perhaps this is in part due to the lack of reliable and ergonomic sampling devices for nasal exhaled air. The only major exception to this is for Fractional exhaled Nitric Oxide (FeNO) measurements, for which there are established protocols and devices for sampling nasally exhaled nitric oxide [21].

In this paper, we have extended the previous studies examining the changes in the levels of volatiles between nasal and oral exhaled breath by exploring a wider range of compounds that are commonly contained in exhaled breath samples [16-18]. Further we have used an analytical technique that affords a higher chemical selectivity than possible with real-time soft chemical ionisation mass spectrometric instruments that were used in the previous studies. We have not only explored the changes in the levels of selected exhaled volatiles between nose inhalation followed by nasal exhalation (nasal) or oral exhalation (oral), but have also investigated a sampling procedure comprised of oral inhalation followed by nasal exhalation, which we will refer to in this paper as the hybrid manoeuvre.

Measurements and comparative analyses are presented to ascertain the extent to which oral microbial activity impact the measured concentrations of the volatile compounds investigated in this study, thereby highlighting the potential limitations of chemically analysing orally exhaled breath samples for the accurate quantification of endogenous trace volatiles of relevance to disease diagnosis and monitoring. We highlight how this issue of recognising bacterial volatile contributions to exhaled breath is becoming ever more important given that machine learning classification algorithms and big data approaches are becoming ubiquitous in breath research.

## 2. Materials and Methods

### 2.1 End-tidal Exhaled Breath and Room Air Sampling

In this observational study, a cohort of twenty-one healthy volunteers with an age range of 20-55 years and a female-to-male ratio of ∼ 1.1:1 was examined. In order to investigate confounders potentially linked to oral microbials, following inhalation through the nose exhaled end-tidal breath samples were collected from each individual via the mouth and via the nose. An additional end-tidal exhaled breath sampling route was performed utilizing a hybrid-manoeuvre consisting of oral inhalation followed by nasal exhalation; notably, this hybrid sampling approach was found to be easier to perform than nasal inhalation-nasal exhalation by all the volunteers. End-tidal breath samples associated with inhalation and exhalation through the mouth were not taken because they do not provide any more information than can be obtained using the three types of exhaled breath sampling investigated in this study. The collection of end-tidal exhaled breath samples, rather than mixed-expired breath samples, was selected because not only are the volatiles contained in these samples generally considered to be close to the alveolar or upper airway concentrations, depending on their solubilities [22], but also because this will maximise the dilution of any volatiles present in the oral cavity as a result of microbial processes. Thus, any differences observed will therefore serve to emphasise more the impact of oral microbial contamination on the breath samples.

Before the breath collection, volunteers were asked to blow their nose (to ensure an unobstructed sampling pathway) and to fill out a consent form. The acceptance criterion required that volunteers must not have eaten or drunk anything (except water) for at least 2 hours. Before each sample collection, volunteers were seated and encouraged to reach a comfortable and calm state. They were instructed to inhale through the nose and exhale through the mouth for at least 3 times before each type of breath sample was collected.

For each volunteer, three clean and sterile glass syringes (Socorex, 250 mL) were utilised for collecting the exhaled breath samples, i.e., one syringe each for (a) oral exhalation (involving nasal inhalation), (b) nasal exhalation (involving nasal inhalation) and (c) hybrid sampling (involving oral inhalation followed by nasal exhalation) for each volunteer. These glass syringes were also used to collect the room air samples, which were taken at each breath sampling collection session and analysed prior to any breath sample measurements to identify potential background room volatile confounders. Before sampling and before any chemical analysis, the glass syringes were stored in an oven at 50 °C for at least 5 minutes, which in the case of the breath samples prevents condensation.

For the exhaled breath samples, each glass syringe was interfaced via a 3-way valve (B. Braun, Discofix, Germany) to a capnometer (NovaMetrix CAPNOGARD 1265 ETCO2 monitor) that was positioned in-line along the sampling route using 1/8^th^ inch perfluoroalkoxy (PFA) tubing. The capnometer was employed to determine visually when the end-tidal phase of the exhaled breath was reached so that sampling could begin (**Figure 1** bottom). Briefly, the 3-way valve was opened in the direction of the glass syringe only when the CO_2_ level in the capnograph reached a plateau (respiratory phase) and the breath sample was manually drawn into the glass syringe. Typically, about 50 mL of end-tidal breath (representing one to two exhaled breaths) were collected into the syringe, which is far more than needed for the analytical device used in volatile identification (see next section) a, which takes 15 minutes per sample, but provides more than an adequate amount of sample if a repeat measurement is considered necessary.

**Figure 1.**
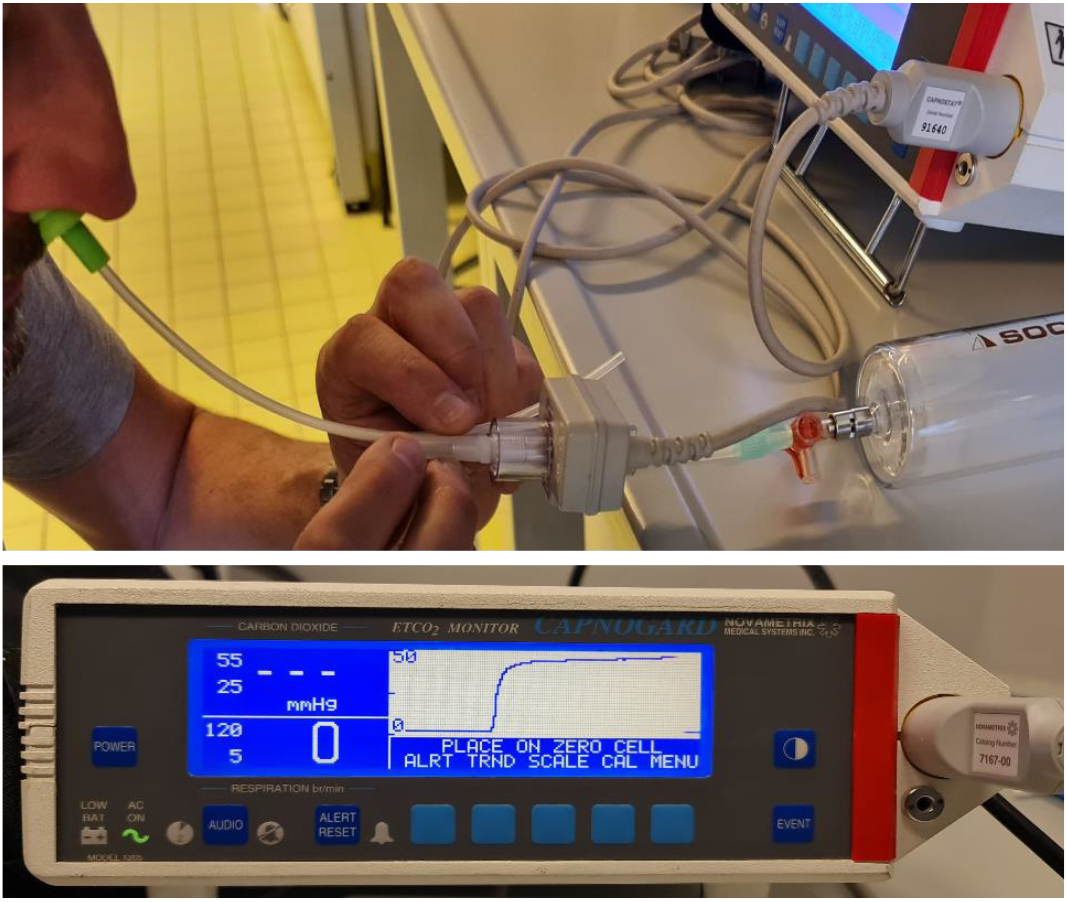
A participant exhaling through one nostril (top) whilst blocking the second nostril and monitoring CO_2_ levels via the capnometer (bottom) to determine when the end-tidal phase of the exhaled breath is reached. As soon as the end-tidal phase of the exhaled breath is reached the valve to the glass syringe is opened and approximately 50 mL of the breath is manually drawn into the syringe. The valve is then closed. The syringe is then removed from the sampling line and placed in an oven at 50 °C until GC-IMS analysis is undertaken. Typically, all samples were measured within 15 minutes of collection.

The in-house made nasal interface consisted of commercially available silicon earplug of the type that are commonly used by swimmers and surfers. Through this a Teflon tube was inserted via a hole drilled along the central axis of the cap; this tube was then connected to a capnometer as illustrated in **Figure 1** (top) via 1/8^th^ inch PFA tubing. No volatile emissions were detected to be emitted from these earplugs. The specially prepared earplug for a given volunteer was inserted into one nostril. The other nostril was pressed closed by the volunteer during the sampling of the exhaled nasal breath.

### 2.2 Breath and Room Sample Analyses using a Gas Chromatograph Ion Mobility Spectrometer

All exhaled breath and room air samples were analysed using a BreathSpec Gas Chromatograph-Ion Mobility Spectrometer (GC-IMS) (G.A.S., Dortmund, Germany [23]). This purpose-built instrument for breath analysis provides the high chemical selectivity of gas chromatography through pre-separation of volatiles in complex sample matrices combined with the high sensitivity of an IMS, with detection limits typically in the low ppb_v_ level with no pre-concentration. The instrument combines an isothermal and flow programmable GC with an IMS. The combination of GC and ion mobility provides an analytical separation in two-time dimensions. The first is according to the GC retention times (in seconds) of the volatiles. The second is associated with the drift times (in milliseconds) of the volatiles’ product ions in the drift region of the IMS.

For this study, the GC was equipped with a 30 m polar chromatographic column (MXT-WAX, L: 30.00 m, ID: 0.53 mm, FT: 0.50 µm), which was maintained at 50 °C. The IMS has a 5 cm drift tube operating at a voltage 2.7 kV (positive ion mode) and also maintained at a temperature of 50 °C. 370 MBq of radioactive tritium (^3^H) is the ionisation source of the IMS through the emission of electrons with a mean energy of 5.7 keV. High-purity nitrogen (5.0, 99.999%) served as both the carrier flow for the GC and the gas flow for the drift tube of the IMS.

We intentionally did not focus our attention on any particular class of compounds in order to replicate the untargeted biomarker discovery scenario. For this reason, a general discovery 15-minute sample analysis pre-set was used. This entailed a fixed flow rate of 100 mL/min for the IMS drift gas, whilst the carrier gas for the GC followed a ramp sequence consisting of a starting flow of 5 mL/min maintained for 30 seconds, followed by a linear increase reaching 50 mL/min after 10 minutes and a final further linear increase up to 75 mL/min reached at the end of the analysis pre-set. These flow settings for the GC allow measurement of compounds up to retention indices of approximately 1470 (GC normalized based on NIST2020 RI Library ‘DB-WAX’).

A breath or room air sample was introduced via an internal pump, passing a 6-port-valve featuring a 1 mL sample loop. For each measurement, and before pressing run on the instrument, 10 mL of the breath or room sample were admitted through a heated (50 °C) sampling line, this being more than sufficient for a high quality BreathSpec measurement and reduces full analysis time. The rest of the sample in the syringe was discarded, because repeat analysis would be too time intensive and prone to sample degradation. After the analysis, the next sample was injected in the BreathSpec. We ensured that no carryover from a previous measurement would affect the following measurement. Following all measurements, syringes were cleaned in hot water containing a detergent, then rinsed with distilled water, and finally placed in an over at 100 °C for six hours prior to their use in any further sampling.

The volatiles identified in the exhaled breath samples by the GC-IMS analysis encompass a diverse range of volatile chemical classes, such as aromatic hydrocarbons, alcohols, ethers, esters, aldehydes, alkanes, alkenes, and ketones.

### 2.3 Data Analyses

The GC-IMS spectra were parsed, processed and analysed using VOCal software (version 0.2.9, G.A.S., Dortmund, Germany [24]), which provides tools for a seamless signal processing/feature extraction/analysis workflow. A qualitative analysis of the volatile compounds was conducted by comparing retention indices and drift times obtained using the GC-IMS results with external libraries (NIST2020 RI) and custom-built databases. Retention indices of volatile compounds were determined using measurements of the retention of homologous 2-ketones C4-C9. IMS drift times are expressed as relative to the position of the reactant ion peak (RIP) as internal standard to compensate for ambient pressure variations. Briefly, peak areas or region of interests (ROIs) were identified on all the spectra simultaneously using a 0.95 percentile criterion. Data were denoised, baseline correction and global cut-off values were adopted.

For each GC-IMS spectrum, the peak volumes (i.e., peak sizes) in arbitrary units corresponding to key volatiles, were calculated and stored in a database for further analysis using Python programming language, namely using the Numpy, Statsmodels, Matplotlib, Seaborn, and Scikit-learn libraries for data parsing, numerical and statistical analysis and visualisation [25–27]. Repeated measures analysis of variance (rmANOVA) was deployed for multiple categorical comparison. Then post-hoc analysis was conducted using multiple pairwise comparisons and considering adjusted p-values to control for the family-wise-error-rate. In case data did not meet the normality and the sphericity conditions, the Friedmann test was used instead of the rmANOVA. The Nyman’s test was then employed to conduct post-hoc analysis following a significant result for a Friedmann test.

Some commonly used data feature extraction strategies were adopted. For example, in the cases where volatile concentrations are sufficiently high that lead to dimer product ion peaks within a measurement set, a linear combination of the monomer product ion (if present) and 2 × the dimer product ion peak volumes was evaluated. Initially, exploratory principal component analysis (PCA [28]) was performed on the whole set of features to mimic typical approaches in untargeted studies, where a lot of data are fed to machine learning and other advanced multivariate algorithms in order to identify patterns in the data. Specifically, before evaluating the PCA, the log_10_ of the peak volumes were first calculated to give less weight to the high-values and reduce the skewness (log peak volume evaluation has proved useful in various other GC-IMS studies [29,30]). Then these data were standardised (using the transformation (x_i_ - µ) / σ) and finally the PCA was conducted.

## 3. Results

**Figure 2**. provides an example of typical GC-IMS spectra recorded for exhaled breath samples collected for an individual following the three different aforementioned breath manoeuvres, namely nasal inhalation / oral exhalation (oral), oral inhalation / nasal exhalation (hybrid), and nasal inhalation / nasal exhalation (nasal).

**Figure 2.**
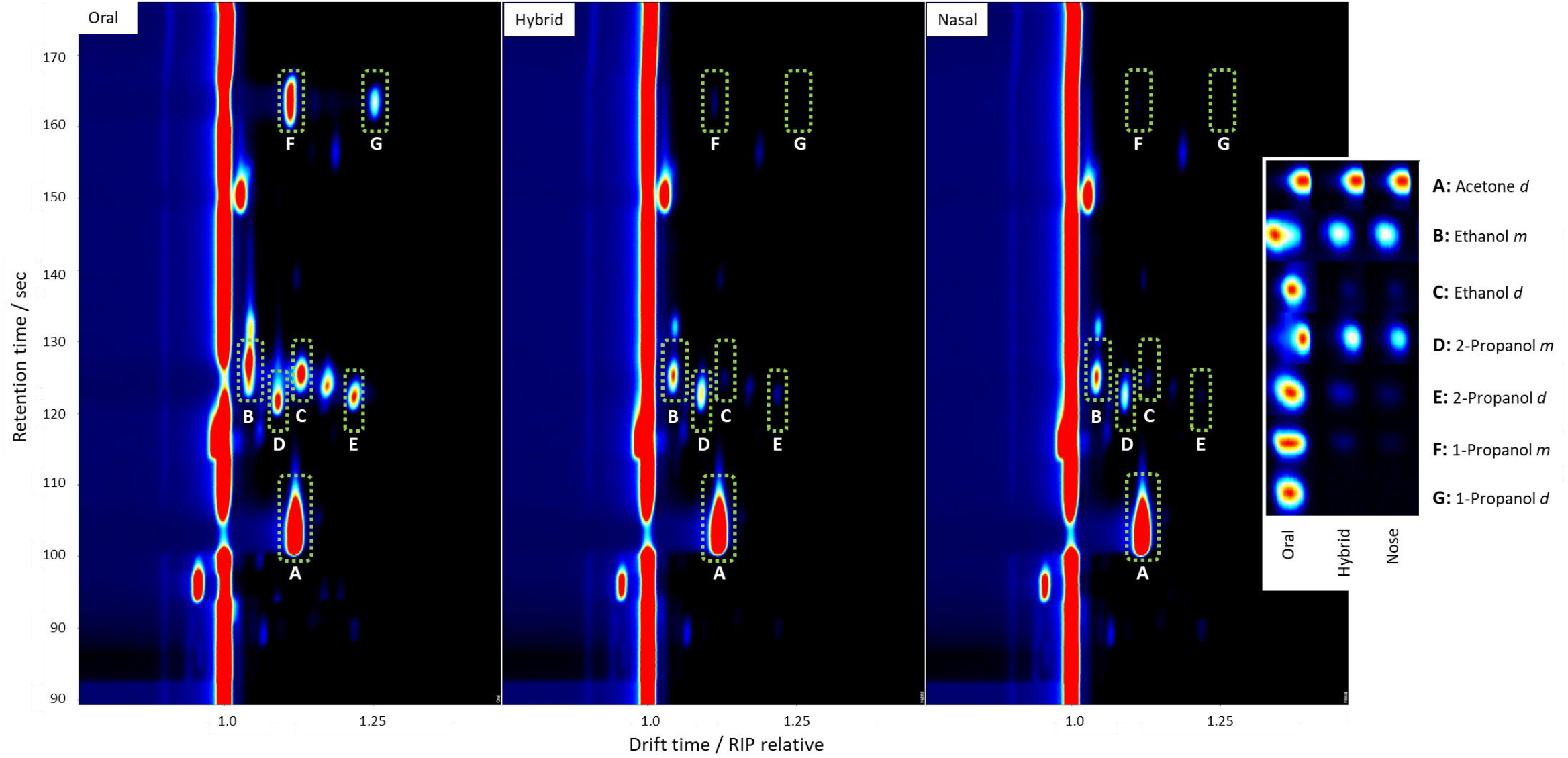
Comparison of typical GC-IMS spectra obtained from the analysis of the three types of breath samples collected from the same healthy volunteer, i.e., nasal inhalation followed by oral exhalation (oral), oral inhalation followed by nasal exhalation (hybrid), and nasal inhalation followed by nasal exhalation (nasal). Each GC-IMS spectrum features the chromatographic volatile retention times in seconds on the y-axis h and the drift times of the product ions on the x-axis, which are shown relative to the reactant ion peak (RIP) at t = 1.0 (i.e., the RIP is the intense red line parallel to the y-axis). The actual drift time of the reagent ions is 7.28 ms. This can be used to determine the real drift times of the product ions using their relative drift times. The RIP intensity is reduced at retention times corresponding to those of the volatiles. This is more obvious for acetone (peak A), which corresponds to the acetone dimer product ion. The acetone monomer product ion has a drift time that is close to that of the RIP and hence is not generally distinguished from the RIP. In these examples, the acetone concentrations in the breath samples are sufficiently high that not only is the RIP completely depleted, but secondary ion-molecule reactions lead to the dimer acetone product ion dominating. Only a small signal associated with the acetone monomer product ion is present. The complete depletion of the RIP means that the acetone dimer peak is saturated. The inset on the right of the three spectra shows the gallery plot view of the key volatiles corresponding to the highlighted peak areas (yellow dashed rectangles). In this inset, the letter *m* stands for the volatile monomer product ion and *d* denotes the volatile dimer product ion.

PCA separated in different clusters, not only breath samples from room air samples, but also oral exhaled air from nasal exhaled air, as shown in **Figure 3**. The loading vectors with the highest magnitude were the ones associated with acetone, methanol, 2-pentanone, isoprene, ethanol, 1-propanol, 2-propanol, and acetoin (3-hydroxy-2-butanone). A further PCA analysis was therefore done on the subset of data corresponding to these selected features. The most significant principal components were the first (percentage of explained variance = 50.79%) and the second (31.99%), which together captured more than 80 % of the variability of the original dataset. Specifically, the vectors linked to acetone, methanol, 2-pentanone, and isoprene are aligned along the separation between breath samples and room air samples (as shown in **Figure 4**). The vectors associated with 1-propanol, 2-propanol, ethanol, and acetoin are linked to the separation between oral exhaled breath and nasal exhaled breath samples, thereby revealing which compounds are truly systemic (released by the blood at the alveolar interface and on the airways surface), which includes in this set of VOCs acetone, 2-pentanone, methanol and isoprene, and those that can largely be generated in the oral cavity, which includes ethanol, 1-propanol, and 2-propanol, and acetoin. The grouped boxplots in **Figure 5** provide an overview of the key volatiles detected in this observational study by comparing the peak volumes in the differently sampled exhaled breath samples and in the room air.

**Figure 3.**
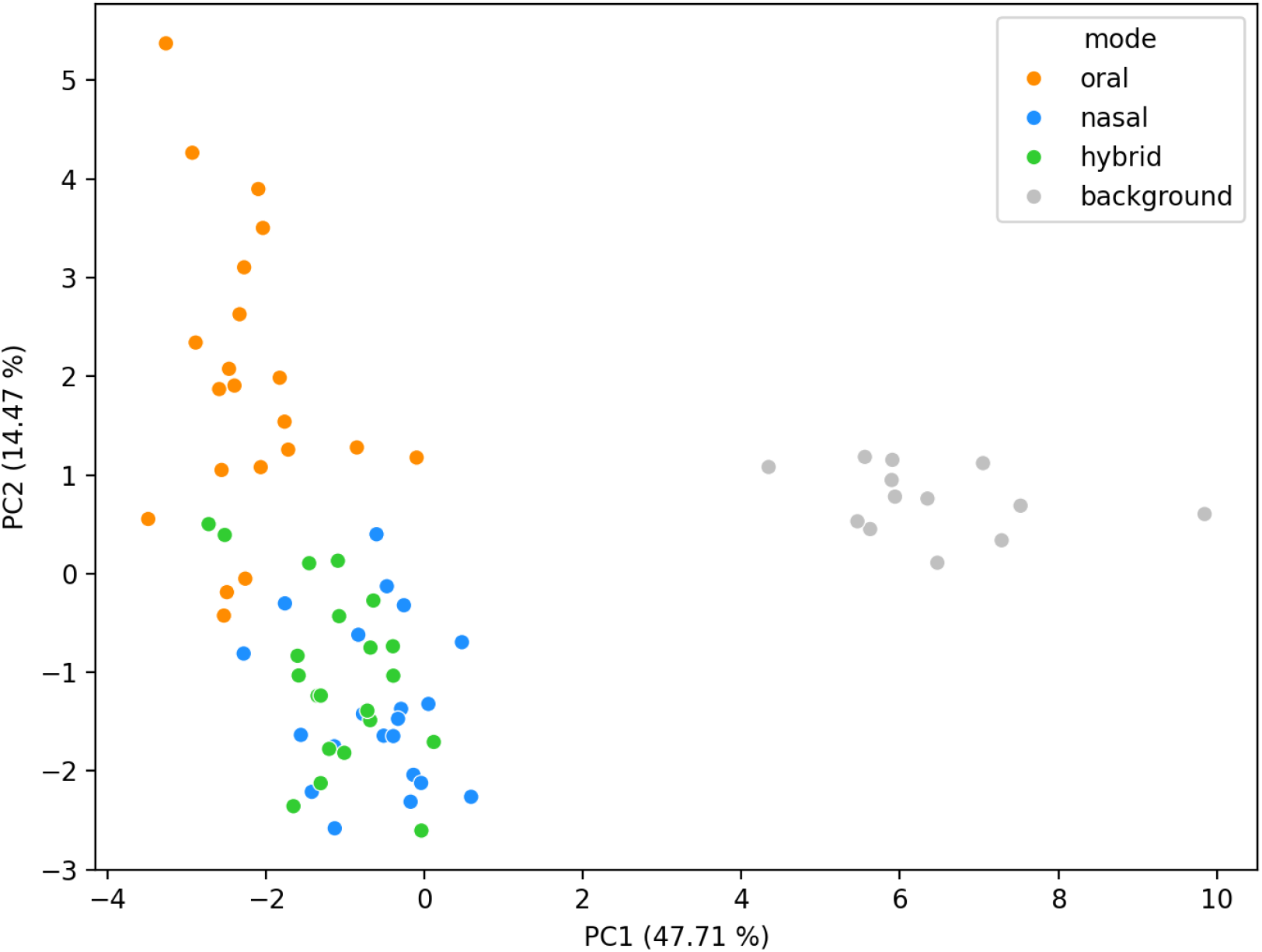
Untargeted Principal Component Analysis of the volatiles found in exhaled breath samples and room air samples (background). The colours indicate the different types of samples. Oral (orange) corresponds to nasal inhalation followed by oral exhalation, nasal (blue) refers to both nasal inhalation and nasal exhalation, and hybrid (green) refers to oral inhalation followed by nasal exhalation. The percentages of the explained variance of the first principal component and of the second principal components are 47.71% and 14.17 %, respectively.

**Figure 4.**
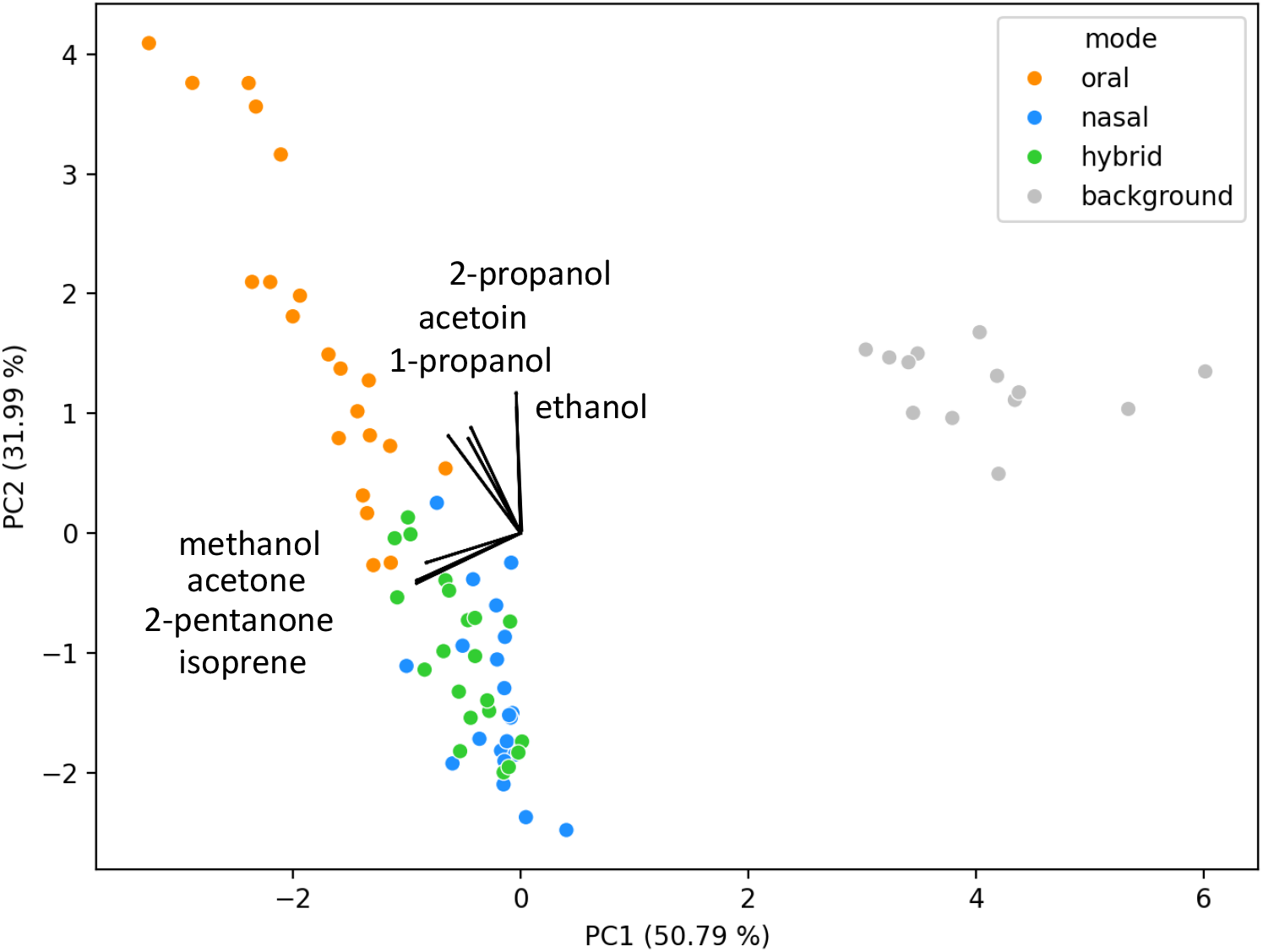
Principal Component Analysis performed after feature selection based on the loadings with the largest values. The percentage of explained variance of the first and second principal components together is more than 80 %. The loadings vectors corresponding to the features acetone, methanol, 2-pentanone, and isoprene are linked to the separation between the breath samples and the background air samples. Whereas the ones corresponding to 1-propanol, 2-propanol, ethanol, and acetoin are associated with the distinction between oral exhaled breath and nasal exhaled breath samples.

**Figure 5.**
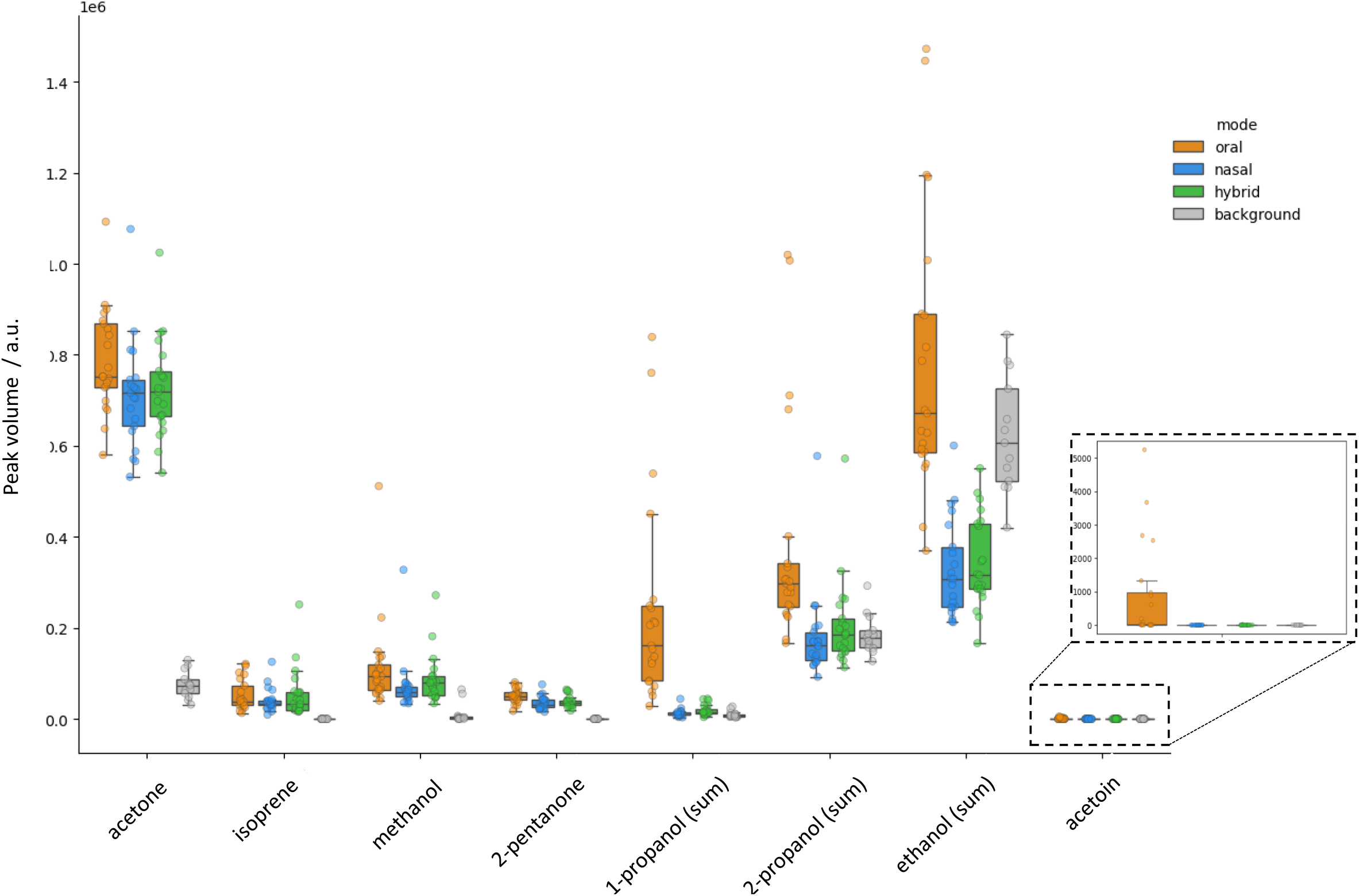
Multi-level boxplot view of the peak volume intensities of the key volatiles selected from the largest loadings of the untargeted PCA analysis. For each volatile, the values of the peak volumes are reported for the three different breath samples and the background air. For those volatiles that result in both monomer and dimer product ions the sum intensities are provided (as discussed in the text). Acetone is of sufficient concentration that only the dimer product ion is observed. For clarity, the inset on the right provides a zoomed-in view of the acetoin peak volume levels in the different sample groups of measurements.

The initial promising results from the PCA analysis highlight the possibility of promptly discriminating volatiles emanating from the oral cavity using GC-IMS on differently sampled breath. This evidence was then corroborated by performing robust comparative statistical analysis to ascertain which specific pairs of groups (e.g., oral versus nasal, or oral versus hybrid) were significantly different.

When comparing different exhaled breath sampling routes, acetone, methanol, 2-pentanone, and isoprene showed similar peak volume intensities (any small difference was found to be statistically insignificant as indicated in **Table 1**), confirming their systemic origin [1,31]. In comparison, ethanol, 1-propanol, 2-propanol, and acetoin have far larger concentrations in the oral exhaled breath samples (see **Table 1**), indicating that these volatiles have strong contributions from the oral cavity. No statistically significant intra-subject difference was detected in VOC concentrations between the two types of nasal exhaled samples, i.e. the ones obtained after inhalation through the nose and the ones collected after inhalation through the mouth (as reported in the Nasal/Hybrid column in **Table 1**.) This provides new and confirmatory data in agreement with complementary studies by Wang et al [14] and Sukul et al [15] with regards to combinations of inhaling either via the nose or mouth and exhaling via the nose.

**Table 1.**
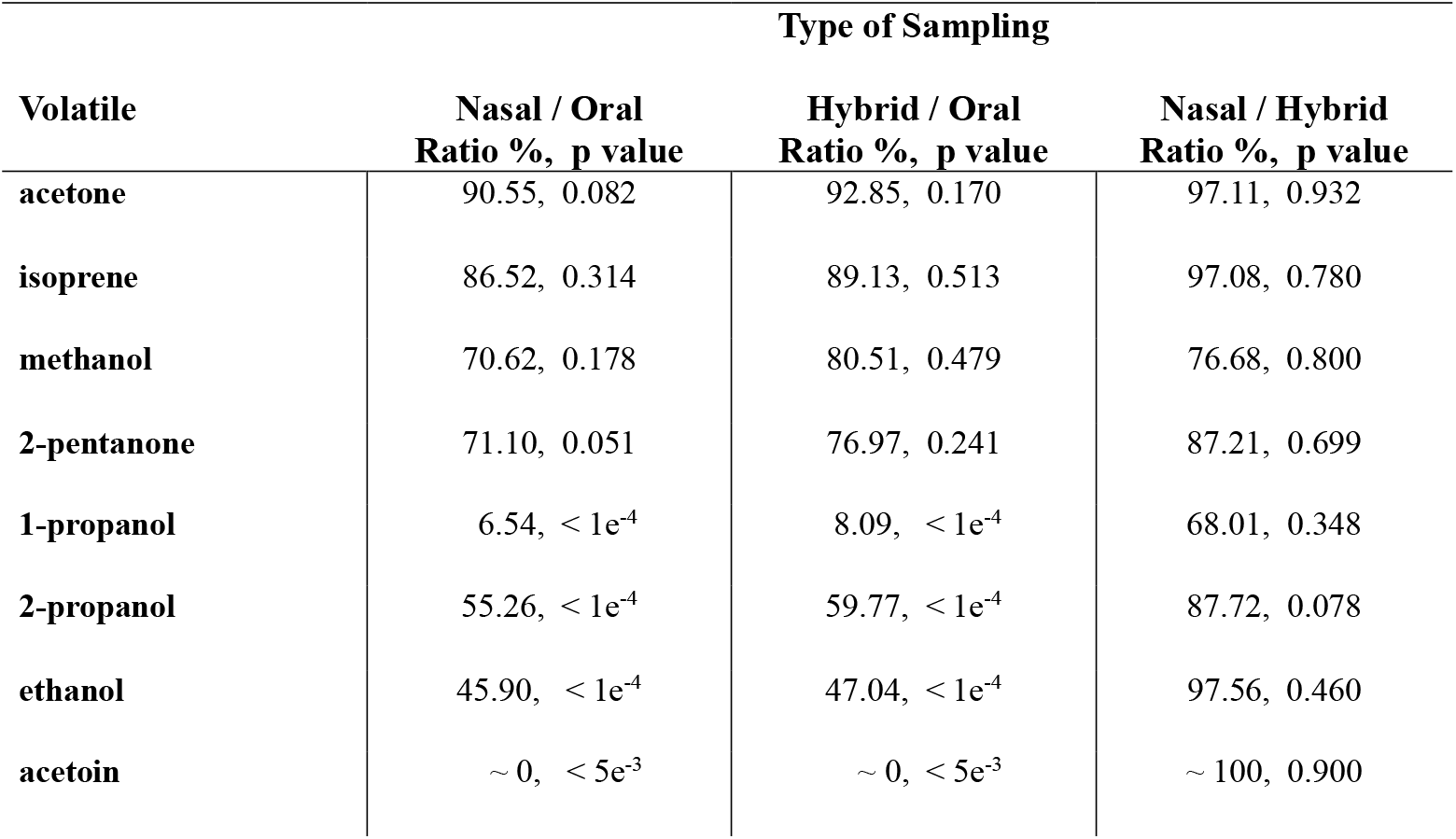
The median ratios (as percentages) of the peak volume intensities for the key exhaled breath volatiles investigated in this study and the adjusted p-values for various statistical comparisons among the different types of sampling.

Inhaled trace concentrations of alcohols are generally considered to be completely absorbed in the upper airways. This can be explained by the airway surface liquid exchange dynamic [32] and by the immune system protection via certain CYPs enzymes present in the lungs [33]. This is in line with our findings, and for ethanol in particular, the results are remarkable: when exhaling through the nose, regardless of the inhalation route, the detected levels of ethanol are far lower than ambient level (see **Figure 6**). Only when exhaling through the mouth, is the intensity of the ethanol higher (background / oral ratio = 90.35%, p-value = 0.11) owing to the microbial activity in the oral cavity.

**Figure 6.**
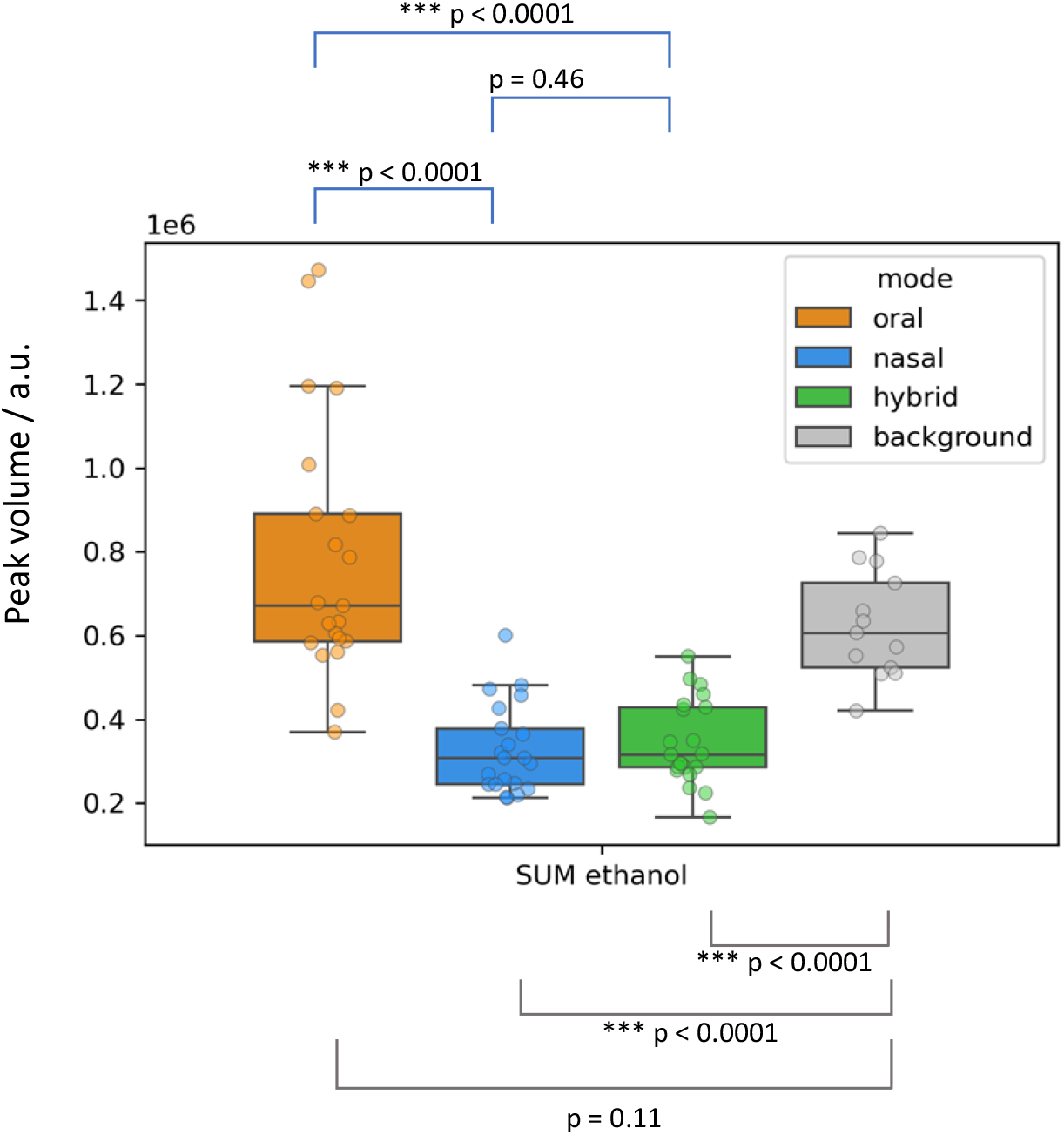
Boxplots for the peak volume intensity of the ethanol detected in the different types of air samples. Nasal exhaled ethanol peak volume levels (either from the purely nasal or the hybrid manoeuvre end-tidal samples) are much lower than the ambient background levels owing to absorption of the inhaled highly soluble ethanol in the upper airways prior to exhalation. This absorption of ethanol in the upper airways is masked in oral exhaled air due to the ethanol being produced in the oral cavity due to the microbiome.

Notably, breath samples of a limited number of volunteers (5 out of 21) featured high levels of dimethyl ether. No significant difference between the intensity of dimethyl ether in oral- and nasal exhaled breath samples was observed. This is considered to be linked to the exposome of those individuals because dimethyl ether is an exogenous substance, commonly used as a propellant gas for cosmetic products that tends to remain in circulation for a long time, and with a long washout timescale from the human body. Exposure factors and for how long certain volatiles remain in the human body provide other challenges in breath research [34].

## 4. Discussion

The application of breath volatile biomarkers for human health needs to establish levels of truly systemic compounds. Previous studies have linked distinct respiratory profiles to various metabolic and inflammatory conditions and attributed to some of these compounds an important diagnostic power [35–37]. However, it is crucial to ascertain that breath biomarker discovery is free from confounding factors, and notably VOCs originating from the mouth microbiome, as these could seriously compromise accurate quantification of volatiles coming from the lungs. This is particularly becoming more of a concern considering how many untargeted studies are conducted by feeding breath sample volatile data to machine learning algorithms or big data and artificial intelligence techniques. Hence, we emphasise that the results of such analyses could be seriously misleading if careful sample preparation is neglected. For example, when attempting to identify infections in the lungs, a machine learning algorithm could pick up elevated levels of acetoin, a compound which is linked to pathogenic bacteria activity [38], as an important classification feature for infections in the airways. However these enhanced levels in oral sampled breath could mask the signal of clinical relevance and lead to biased results and false positives.

The results presented in this study indicate that oral microbial activity is responsible for marked variations between orally and nasally sampled exhaled breath for certain volatiles, allowing for the systemic compounds to be identified with a higher level of confidence. It is worth mentioning that mechanical filter ciliated body and upregulation of iNOS and nitric oxide keep the nasal environment with a minimal microbial load [39], thus making the nasal route a the preferred choice for collecting higher quality exhaled breath samples that are little contaminated by microbial volatiles. Interestingly, the non-significant difference in VOC concentrations between the two types of nasally exhaled breath samples (inhalation through the nose or through the mouth) gives confidence that inhaling air through the mouth does not significantly affect the concentration of the end-tidal exhaled volatiles owing to the loss of volatiles with high solubility in the upper airways.

Whilst oral sampling is more practical with current commercially available devices, our findings indicate that collecting nasal exhaled breath samples, and preferentially end-tidal samples, for spectroscopic, spectrometric or mass spectrometric analyses are preferred compared to oral exhaled breath samples owing to the considerable reduction in confounding volatiles relating to microbial activity. We recommend that, where possible, untargeted clinical studies should focus on nasal exhaled end-tidal breath collection independent of the inhalation route. Manufacturers could facilitate this by developing and providing ergonomically and simple-to-use nasal interfaces with CO_2_ detection capabilities that can address this crucial challenge in the field of breath research. Nasal interfaces should be as comfortable and as non-invasive as possible, especially considering that the elderly, children and critically ill patients might find nasal sampling uncomfortable. In terms of comfort, we explored a hybrid-breathing manoeuvre consisting of inhalation through the mouth (whilst comfortably wearing the nasal interface) and exhalation through the nose, which proved to be easier to perform for all the volunteers in this study than nasal inhalation-nasal exhalation.

## 5. Conclusions and Outlook

It is often stated that more standardisation is needed in breath research [40–42], and that it is imperative to identify confounders and establish sampling procedures that can avoid them. Further, extensive research is necessary to determine the origin of most exhaled VOCs in order to help make breath analysis a valuable tool for clinical disease diagnosis. The work reported in this paper highlights how oral microbial activity is a serious confounding concern when it comes to the use of orally sampled exhaled breath to identify volatile biomarkers for medical diagnostics in untargeted breath research studies.

Specifically, we investigated the intensities for a number of volatiles in exhaled breath measured with GC-IMS, comparing the differences in breath samples between nasal inhalation followed by either oral or nasal exhalation or a hybrid sampling involving oral inhalation followed by nasal exhalation for a cohort of twenty-one volunteers. Significant rises in the concentrations of volatiles in oral compared to nasal exhaled breath samples provide information on those volatiles that are predominantly produced in the oral cavity, as has been suggested in other previous studies [14,16,17].

Unfortunately, the effects of microbial activity on the exhaled breath volatilome are often neglected in breath research studies, where it is crucial to identify variations in the concentration of compounds contained in exhaled breath to metabolic and physiological processes. VOCs produced from microbials in the mouth can mask the levels within the systemic circulation, thereby jeopardising any attempt to pursue clinical application for monitoring/diagnosing disease. Owing to the unique nasal cavity properties (robust immune defence system, mucus, hair and nitric-oxide rich environment) causing minimal microbial load in the nose, an exhaled breath nasal sampling approach is less affected by non-systemic volatile confounders, thereby holding better promise for untargeted breath biomarker discovery.

Given that previous studies investigating nasal versus oral exhaled breath samples appear to be overlooked, we hope that this work will draw the attention of breath researchers to consider adopting nasal sampling as the preferred and more robust route for improved measurements, thereby ensuring better quality assured and controlled results. The variability in breath sampling methodologies and the resulting challenges in data comparison and integration are significant contributing factors. To help realise the full potential of breath analysis for use in clinical and pre-clinical medicine, the standardisation of breath sampling and analysis techniques is essential.

The major outcome of this study is that it highlights the need to critically review breath research findings and focus on developing a new generation of practical and economically viable nasal interfaces to provide more accurate end-tidal exhaled breath sampling. A secondary important outcome of this study is the observation that when examining the hybrid (oral inhalation before nasal exhalation) breath sampling manoeuvre, there is no statistically significant difference compared to the purely nasal manoeuvre. Not only is this reassuring, but it also holds great promise in terms of developing ergonomic ways to collect nasal breath sample in an easy and acceptable way: the nasal probe can be comfortably positioned prior to the collection of the nasal exhaled breath sample, rather than quickly attempting to exhale through the nasal cannula immediately after a free nasal inhalation, thereby mitigating the need for a complex nasal sampling device incorporating one-way valves.

## Ethics

This study was approved by the Institutional Ethics Committee of the Medical University Innsbruck. All volunteers were explained the significance of the study and had to give written Informed Consent prior to providing a breath sample. The study protocol conforms to the ethical guidelines of the 1975 Declaration of Helsinki.

## Data availability statement

All data used in this publication are available upon request from the corresponding author.

## Acknowledgments

We acknowledge the EU HORIZON Innovation Actions HORIZON-CL3-202-DRS-01-05, Project Number 101073924 (ONELAB) for funding Dr. Lorenzo Petralia’s postdoctoral position. Anesu Chawaguta acknowledges the FFG IKT der Zukunft, IKT der Zukunft, IKT der Zukunft-Resilienz und Distancing (DEVICE) for funding his PhD position.

